# Spatiotemporal-multimodal integration reveals BCG-induced skin-blood crosstalk

**DOI:** 10.1101/2025.11.19.689184

**Authors:** Amrit Singh, Ole Bæk, Frederik Schaltz-Buchholzer, James Campbell, Nicholas P West, Basam Elgamoudi, Anita J. Campbell, Elsi Cá, Scott J. Tebbutt, Casey P. Shannon, Peter Aaby, Tobias R. Kollmann, Christine Stabell Benn, Nelly Amenyogbe

## Abstract

Tuberculosis (TB) remains the leading cause of infectious death. The Bacille Calmette-Guérin (BCG) vaccine has been the only licensed vaccine available for TB prevention. Despite BCG being administered intradermally for over a century to >100 million individuals annually, the molecular events in the skin following BCG administration have not been investigated; as a result, measurable correlates of protection that could predict vaccine effectiveness already early after vaccination are lacking. Here we show that BCG immediately (within one day after vaccination) induces dynamic molecular waves that drive the acute human host response across space (layers of the skin and systemically in blood) and time (days). Integration of this data across space and time identified robust networks of interactive modules related to immune surveillance (e.g. Langerhans cells), cell trafficking (e.g. endothelial cells, *ITGB5*), and trained immunity (e.g. neutrophils, macrophages, γδ-T cell). Importantly, not only were we able to identify BCG-activated pathways associated with ‘trained immunity’ such as mTOR signaling and glycolysis/gluconeogenesis, we were able to pinpoint the time-point and precise location (skin layer) of the initial activation of theses pathways. Combining tissue biopsies of human skin (spatial genomics) with ‘liquid biopsies’ (cell-free blood plasma RNASeq) following BCG vaccination our data both confirmed known evidence (e.g. prominent γδ-T cell induction at the site of BCG administration; negative correlation of blood vs tissue myeloid-derived suppressor cells), but also generated promising new leads such as baseline levels of B cells, platelets and nuocytes in the skin prior to BCG administration predict eventual outcome, and that these predictive differences in baseline cellular composition can be captured non-invasively using high resolution images of the site of injection (dermatoscopy). Given this data represents the first holistic view of the acute molecular response to BCG in the skin in a human population at medium to high TB risk, we anticipate our findings of the immediate/early events following BCG vaccination, including non-invasive predictive assessment will support acceleration of vaccine development in the fight against TB.

## Introduction

Tuberculosis (TB) remains one of the most pressing global health challenges, causing significant morbidity and mortality in millions of people each year. Despite efforts to control and prevent its spread, TB continues to be the leading cause of infectious death^1^. The Bacille Calmette-Guérin (BCG) vaccine is currently the only licensed vaccine available for TB prevention and is administered to >100 million individuals annually. However, the efficacy of BCG vaccination varies dramatically across different populations^2^. The mechanisms behind this variation in BCG vaccine efficacy are unknown, as are measurable correlates of protection that could predict vaccine effectiveness early in clinical trials to accelerate progress in improving TB control.

BCG is currently given as an intradermal injection into the epidermis resulting in a blanched “wheal” of ∼5mm in diameter (∼10mm for the adult dose) that quickly dissipates^2^. Over the subsequent days an inflammatory response is initiated, which over weeks to months develops into a pustule, eventually forming a 5-7mm scar. This scar is a visible indicator of the local immune response to the vaccine and is traditionally considered evidence of successful “vaccine take”^2^. The local response in the skin leading to clearance of BCG has been shown to correlate with systemic markers of TB protection (for example, in mycobacterial growth inhibition assays), and represents a human controlled infection model (CHIM)^3–5^. Even though the BCG vaccine recently celebrated its 100^th^ birthday and has been administered to *billions* of people, the molecular and cellular events locally in the skin upon intradermal BCG vaccination have never been fully investigated. Absence of insight into this initial host response to BCG in the skin have contributed to the lack of systemic biomarkers in peripheral blood, which could predict host immune responses.

We hypothesized that BCG vaccination induces early molecular changes in the skin that predict eventual scar formation and that these initial local events can also be detected systemically through blood-based biomarkers. Blood-based systemic assessment (vs tissue biopsy) could serve as a “liquid biopsy,” offering an accessible, minimally-invasive tool to track immediate local immune responses to BCG. Such ‘tissue biomarker’ approach could accelerate the effort to determine how and when BCG confers protection against TB, facilitating the prediction of vaccine efficacy in larger vaccine studies.

To test our hypothesis, we performed a comprehensive, longitudinal randomized experiment involving individuals who did not have a BCG vaccine scar. Following administration of BCG, we collected skin and blood samples at different time points allowing us to capture the dynamic changes occurring in both the skin and systemic circulation over time. We paired these data with dermatoscopic images at the site of inoculation to monitor the progression of the developing skin reaction. Skin biopsies were analyzed using spatial multiomics, including identification of cell-specific markers as well as whole-transcriptome spatial gene expression profiling. We also assessed changes in cell-free RNA in peripheral blood plasma samples as “liquid biopsies” capturing early local molecular changes that can be detected systemically.

## Results

### BCG can be detected in the skin during the first week post vaccination

In this randomized experimental study in Guinea-Bissau, we enrolled 19 adult, non-pregnant HIV-negative females without signs of active tuberculosis (Fig.1A). Four females were randomized to the placebo group whereas 15 were randomized to receive the BCG vaccine. Baseline blood samples were collected prior to placebo/BCG vaccine on day 0, and participants were randomized for one additional blood draw and a skin biopsy collected either on day 1, 7 or 14 post-enrolment (Fig.1B). Dermatoscopic images of the site of BCG administration were obtained for all participants at baseline (immediately following vaccination) to visualize the post-injection wheal and at follow-up prior to biopsy sample collection, to visualize the developing BCG skin reaction. In blood samples from baseline and follow-up, cell-free plasma was obtained and cell-free RNA sequencing was performed. Skin biopsies obtained during follow-up were both stained and spatially-profiled for whole transcriptome gene-expression across multiple skin regions (epidermis, dermis and hypodermis, Fig.1A).

**Figure 1.**
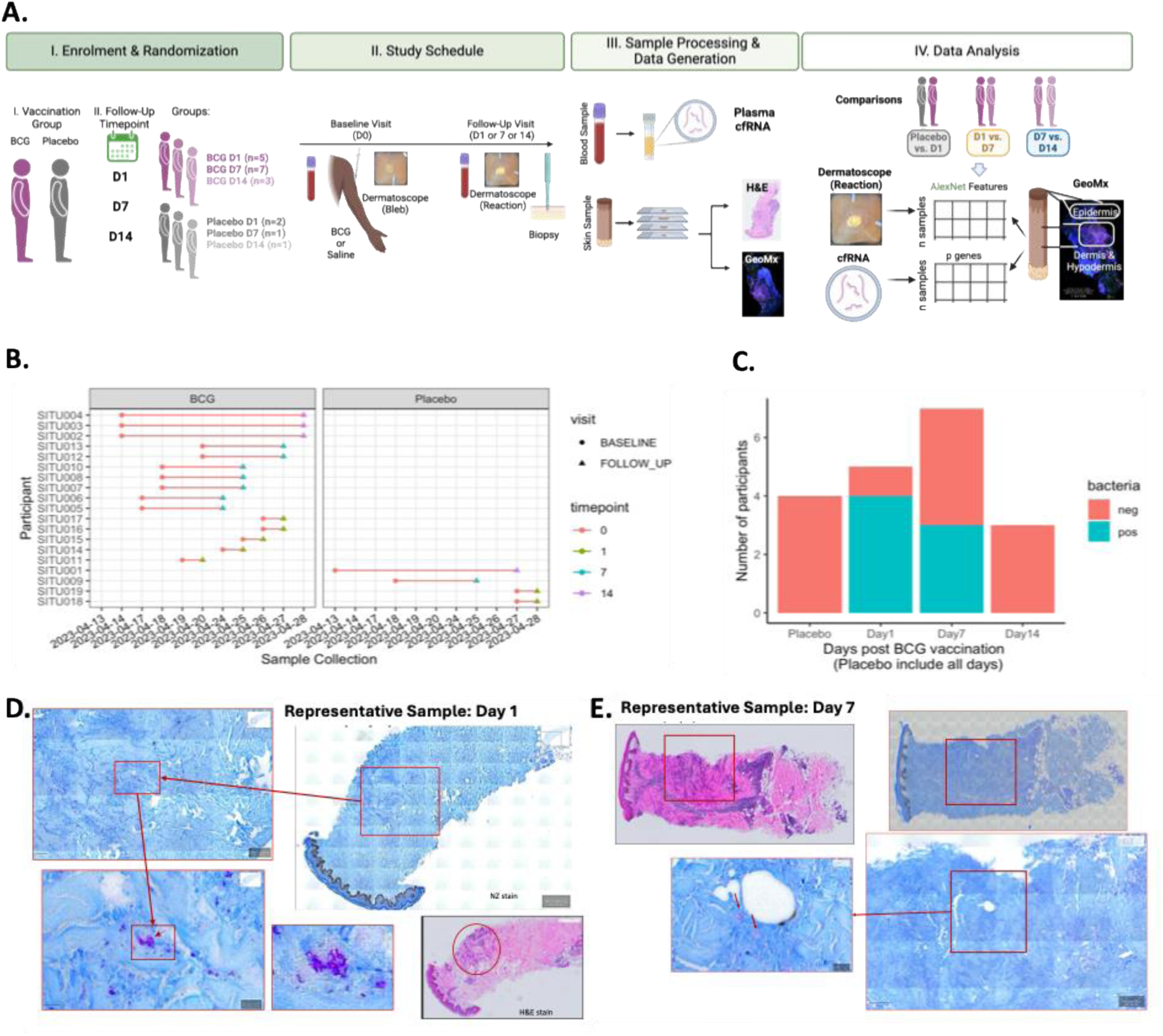
Bacille Calmette-Guérin is detectable in areas of cellular infiltration. A. Study design, sample processing, and data analysis strategy. B. Sample collection dates of participants including baseline and follow-up samples at days 1, 7 and 14 C. Number of participants with skin biopsies in which Mycobacteria was detected. D. H&E and ZN-stained skin samples obtained one day after BCG vaccination. E. ZN and H&E staining of skin samples obtained during the day 7 post-BCG vaccination.

BCG or placebo was successfully administered intradermally to all participants as assessed by the dermatoscopic images of the injection site (Table S1). Dermatoscopic images show a progression of the inflammatory response over time, where established ulceration, post inflammatory hyperpigmentation and reduction in erythema can be observed at day 14. The images at Day 7 displayed heterogeneity, with some participants already showing development of a pustule, scale and early ulceration while others only showed redness and swelling. Some images from day 7 depicted rosettes typically related to scale or fibrosis around adnexal structures and openings that can be found in inflammatory and infectious conditions, skin cancers and actinic keratoses. ^6,7^. This indicates that individual variation in the local immune response can be captured early and non-invasively through dermatoscopic images.

Ziehl-Neelsen (ZN) staining identified the presence of mycobacteria in skin biopsies obtained at days 1 and 7 but not on day 14 post-BCG vaccination and not at all among placebo recipients (Fig 1C). BCG vaccination led to rapid development of cellular infiltrates on Haematoxylin-Eosin (HE) and ZN stains, already detectable from Day 1 and onwards (Fig 1D-E). The dermatoscopically visible inflammatory response and development of the pustule continued after mycobacteria were no longer detectable at the site of injection. Thus, clear signs of both macro-and microscopic inflammation could be identified in all BCG-vaccinated participants but not in controls.

### BCG induces distinct transcriptomic responses in different layers of the skin

To profile the development of the local host response to BCG, skin biopsies were sectioned with regions from the epidermis, dermis and hypodermis profiled for 18,677 gene-transcripts using spatial transcriptomics. Principal component analysis showed progressive differences in the transcriptome between placebo vs BCG over time in the epidermis, dermis and hypodermis (Fig.S1). Given the limited number of hypodermis regions that were profiled, the dermal and hypodermal data were pooled and analyzed together (dermis+hypodermis). To investigate the dynamic changes in the local skin response, we performed differential gene-expression analysis comparing placebo with Day 1, Day 1 with Day 7 and Day 7 with Day 14 post-BCG vaccination. The strongest signal was observed when comparing Day 1 post-BCG with placebo in both the epidermis and dermis+hypodermis (Fig. 2A). At a Benjamini-Hochberg False Discovery Rate (FDR) of 10%, 4,531 and 1,704 differentially expressed gene-transcripts were identified when comparing placebo with day 1 post-BCG samples in the epidermis and dermis+hypodermis, respectively. At the same FDR, only 29 gene-transcripts were differentially expressed between day 1 and day 7 post-BCG vaccination. In the epidermis, 1,428 and 3,103 gene-transcripts were up and down-regulated, respectively, whereas in the dermis+hypodermis 1,171 and 533 gene-transcripts were up and down-regulated one day after BCG vaccination, when compared to placebo samples (Fig.2B). Although most differentially expressed genes did not overlap between the two skin regions, 660 and 280 gene-transcripts were up and down-regulated in both regions (epidermis and dermis+hypodermis, Fig.2C). All of the 29 differentially expressed genes between day 1 and day 7 were up-regulated at day 7 as compared to day 1 post-BCG in the epidermis (Fig. 2C). Interestingly, 16 genes of these gene-transcripts were down-regulated at day 1 as compared to placebo, but were up-regulated at day 7 post-BCG vaccination. Geneset enrichment analysis was then performed using genesets from the Kyoto Encyclopedia of Genes and Genomes (KEGG) database. Fig.2D depicts the normalized enrichment scores each for 70 significant genesets (FDR<1%) across comparisons and skin tissues. In general, various cellular processes and functions were activated throughout the first two weeks after BCG vaccination (see genetic information processing and cellular processes, Fig. 2D). These included genesets related to folding, sorting and degradation (proteasome, ubiquitin mediated proteolysis, protein export, RNA degradation) plus glycan biosynthesis and metabolism (cell cycle and apoptosis). Genesets related to the immune system (*e.g.,* antigen processing and presentation, leukocyte transendothelial migration, toll-like receptor signaling pathway) were activated at day 1 post-BCG vaccination in both the epidermis and dermis+hypodermis, but only the dermis+hypodermis at day 14 (see organismal systems in Fig. 2D). Genesets related to signal transduction (*e.g.,* VEGF signaling pathway, mTOR signaling pathway, TGF-beta signaling pathway) were activated only at day 14 in the dermal layers (see environmental information processing in Fig. 2D). The cytokine-cytokine receptor interaction pathway was activated at day 1 but suppressed at day 7 and day 14 in both the epidermis and dermis+hypodermis. Interestingly, olfactory transduction and neuroactive ligand-receptor interaction were suppressed throughout the entire two-weeks post BCG-vaccination.

**Figure 2.**
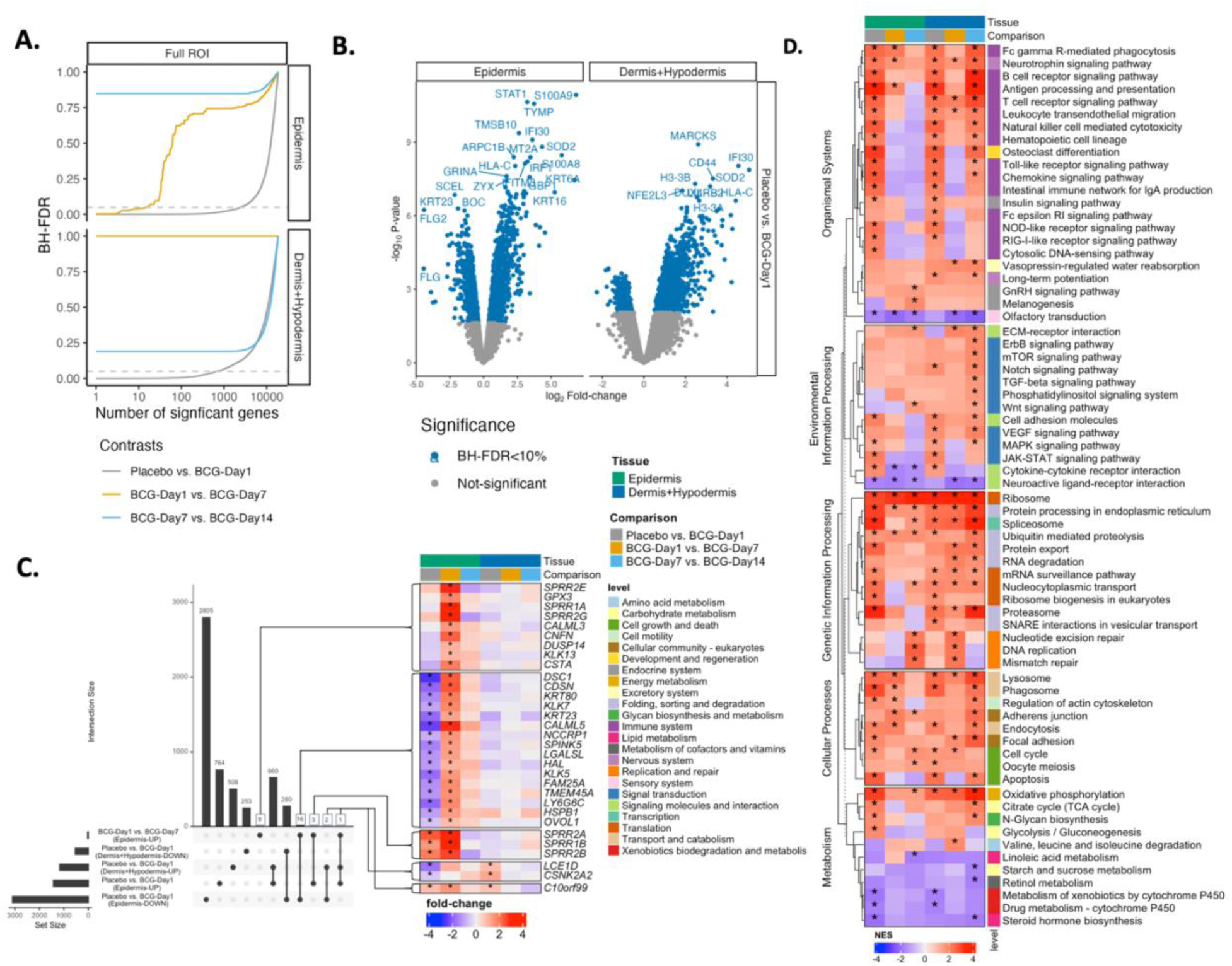
Spatial gene-expression analysis in the skin comparing placebo and BCG samples over time. A. Differentially expressed genes across various comparisons and tissue-regions and Benjamini-Hochberg False Discovery Rate (BH-FDR) thresholds. The dashed line represents a BH-FDR cut-off of 10%. B. Volcano plot depicting the placebo vs. day 1 post-BCG comparison results for the epidermis and dermis+hypodermis regions. Blue dots indicate significant gene-transcripts (BH-FDR<10%). C. Overlap between the significant gene-transcripts identified across all comparisons in A (epidermis: placebo vs. day 1, dermis+hypodermis: placebo vs. day1 and epidermis: day1 vs. day 7). The heatmap depicts the log_2_ fold-change between the pairwise comparisons (placebo vs. day 1, day 1 vs. day 7 and day 7 vs. day 14) for significant genes in multiple comparisons and the epidermis, where the color indicates up/down regulation. For example, for the placebo vs. day 1 comparison, red (purple) depicts up (down)-regulated gene-transcripts at day 1 post-BCG vaccination compared to placebo samples. D. Heatmap of the significant KEGG pathways (BH-FDR<1%). A positive (red) NES depicts up-regulation of a geneset (*e.g.* pathway) in a given comparison and tissue-region whereas negative (purple) depicts down-regulation of a geneset in a given comparison and tissue-region. Stars indicate significant genes/genesets.

### Distinct transcriptional responses to BCG in CD3+, CD31+ and CD68+ T Cells

Geneset enrichment analysis was also performed using cell-type marker genes from the Panglao database^8^ (Fig.3A). Given that these findings are based on cell-type marker genes, an increase/decrease for a given cell-type is based on the increase/decrease in the expression of cell-marker genes (*e.g.,* representative fold-changes of neutrophil markers genes are shown to be increased at day 1 and decreased in activity at day 14 in the epidermis). Cells of the immune system such as natural killer T cells, dendritic cells, macrophages and neutrophils were up-regulated at day 1 in both the epidermis and dermis+hypodermis. However, all of these cells had decreased in the epidermis by day 14 as compared to day 7 yet remained elevated in the dermis+hypodermis. γδ T cell responses increased throughout the two-week period post-BCG vaccination in both the epidermis and dermis+hypodermis, while fibroblast cell activation was observed only at the later time points and across all regions of the skin.

**Figure 3.**
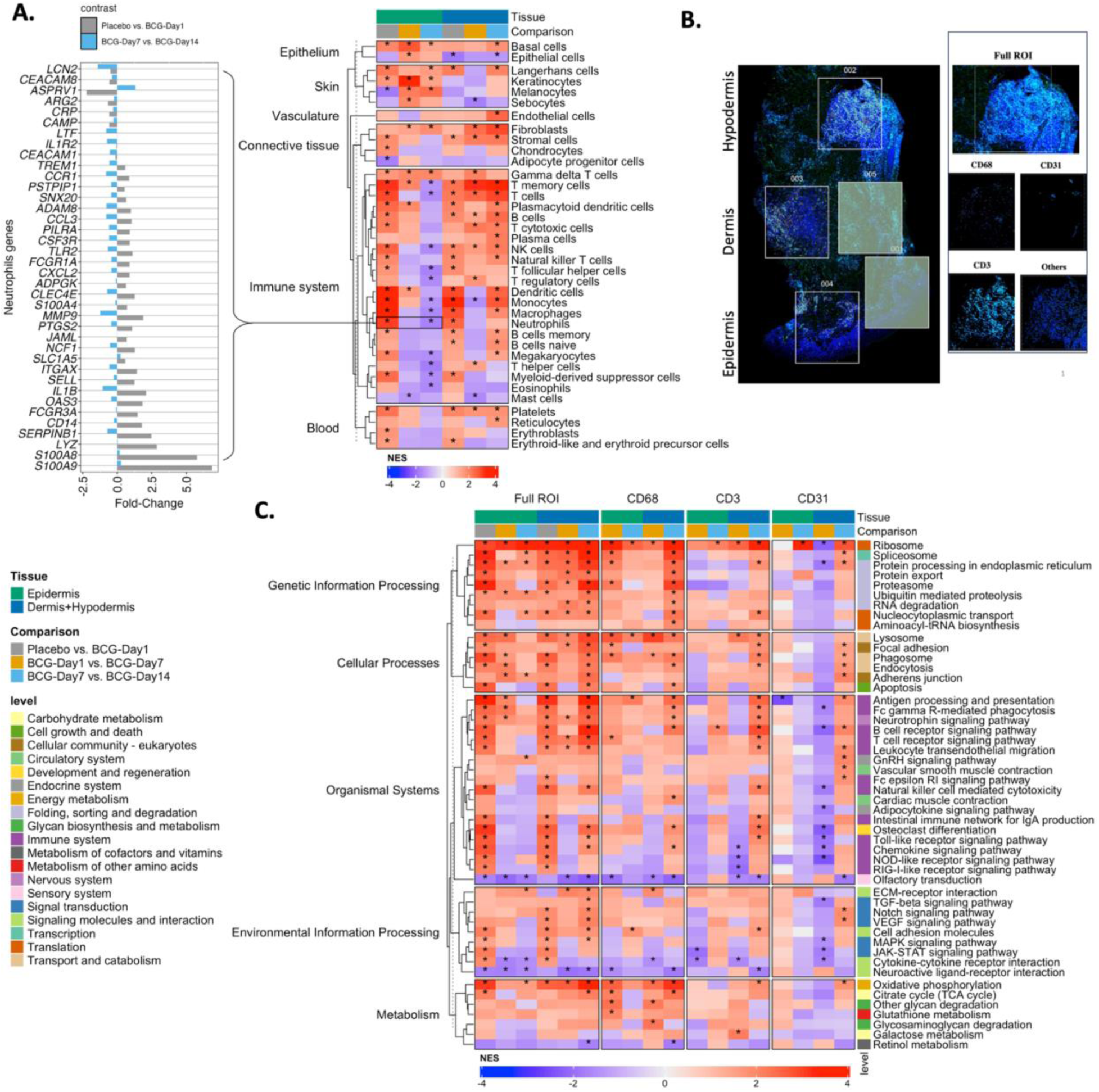
Spatial cell-specific gene-expression analysis in the skin comparing placebo and BCG samples over time. A. Heatmap of Normalized Enrichment Scores (NES) from gene set enrichment analysis (GSEA) of cell-types based on three pairwise comparisons (placebo vs. day 1, day 1 vs. day 7 and day 7 vs. day 14) restricting to regions from the epidermis or dermis+hypodermis. A positive (red) NES depicts up-regulation of a cell-type in a given comparison and tissue-region whereas negative (purple) depicts down-regulation of a cell-type in a given comparison and tissue-region. Stars indicate significant cell-types based on a BH-FDR of 1%. B. Digitized immunofluorescence image of a representative skin section highlighting the regions of interest (ROI) selected for gene-expression profiling C. NES from GSEA of pairwise comparisons using the Full ROI (similar to bulk expression) or segmented by cell marker for different tissue regions in the skin.

To further delineate these responses, we segmented the spatial regions (Fig.3B) based on CD3+ (T cells), CD31+ (endothelial cell) and CD68+ (phagocytic cell) staining to allow for cell-specific differential gene-expression analysis (day 1 vs. day 7 and day 7 vs. day 14). At an FDR of 1%, 32, 31 and 54 significant KEGG pathways were identified for CD3+, CD31+ and CD68+ cells, respectively. Fig.3C depicts the Normalized Enrichment Scores (NES) for these gene sets from each cell-type as well as the bulk data (Full ROI) from Fig.2D. Phagocytic (CD68+) cells exhibited activation (increased activity) of several key pathways (*ribosome, lysosome, focal adhesion, antigen processing and presentation, oxidative phosphorylation*) during the two weeks post-vaccination regardless of tissue region. Interestingly, endothelial cells (CD31+) showed little to no initial transcriptional response in the epidermis but displayed significant down-regulation of pathways at Day 7 post-BCG vaccination in the dermis+hypodermis layers, a trend that was reversed by day 14 (*e.g., ribosomal activity, protein processing in endoplasmic reticulum, B cell receptor signaling pathway*). Many pathways such as the *chemokine signaling pathway* and *cytokine-cytokine receptor signaling* were significantly down-regulated in CD3+T cells at day 7 as compared to day 1. Interestingly, *antigen processing and presentation* was up-regulated in CD68+ phagocytic cells but down-regulated in CD31+ endothelial cells in the epidermis.

These findings suggest that BCG vaccination induces a complex, time- and location-dependent host response in the skin involving several cell types and pathways. Overall, the immediate upregulation of immune pathways indicates that BCG vaccination rapidly activated both innate and adaptive immune responses in the skin. The persistence of this immune activation in the deeper skin layers (dermis+hypodermis) may be crucial for the development of long-term immunity and later scar formation. Since most of the observed responses can be linked to systemic immune functions, these local skin changes might reflect or contribute to systemic immunity against TB that is elicited by the vaccine.

### BCG induces a pronounced cell-free transcriptional response in plasma, reaching full bloom later than skin-based changes

To better understand systemic effects of the BCG vaccine, we sequenced RNA transcripts in cell-free plasma as a systemic (peripheral blood) response that captures local (skin) events as a “liquid biopsy” on blood samples collected at baseline and all follow-up time points. Principal component analysis applied to the resulting dataset of 19,907 gene-transcripts depicted a clear separation between baseline and day 1 and day 7 samples for the BCG group only (Fig.S2). Gene-expression data for each participant was normalized to baseline levels (day 14 minus day 0 and day 7 minus day 0); and as for the analysis of skin transcripts, three pairwise comparisons were made: placebo vs. day 1, day 1 vs. day 7 and day 7 vs. day 14. While univariate analysis did not identify significant gene-transcripts at a FDR of 10%, geneset enrichment analysis (GSEA) identified 2 KEGG pathways for the placebo vs. day 1 comparison and 35 KEGG pathways for the day 7 and day 14 comparison (Fig.4A). With the exception of 8 pathways related to metabolism (*e.g., tyrosine, ether lipid and tryptophan metabolism*), all other pathways were up-regulated at day 14 as compared at day 7. Up-regulated pathways during the second week post-BCG vaccination included various immune pathways such as *toll-like receptor, RIG-I-like receptor, T cell receptor, B cell and chemokine signaling pathways*. At an FDR of 10%, GSEA also identified 8 types of immune cell markers that were up-regulated at day 14 as compared to day 7 (Fig.4B). These included both phagocytic cells (monocytes, macrophages, plasmacytoid dendritic cells), megakaryocytes and various T cells (natural killer, and γδ T) cells. In addition, exploratory analysis using sparse partial least-squares (sPLS) captured an association between baseline levels and changes of features in dermatoscopic images relative to baseline (Fig.4C). Next, sPLS was used to regress 1,938 dermatoscopic image features (post vaccination minus baseline) onto baseline levels of 29 cell-types (estimated from plasma cfRNA) from 15 BCG-vaccinated participants. Fig.4C depicts the samples grouped based on the day post-BCG vaccination along Component 2. Interestingly, higher baseline levels of B cells and platelets were observed in samples from participants who rapidly developed a noticeable scar, whereas higher levels of nuocytes^9^, innate effector cells related to Type 2 responses, were observed in samples from participants who did not.

**Figure 4.**
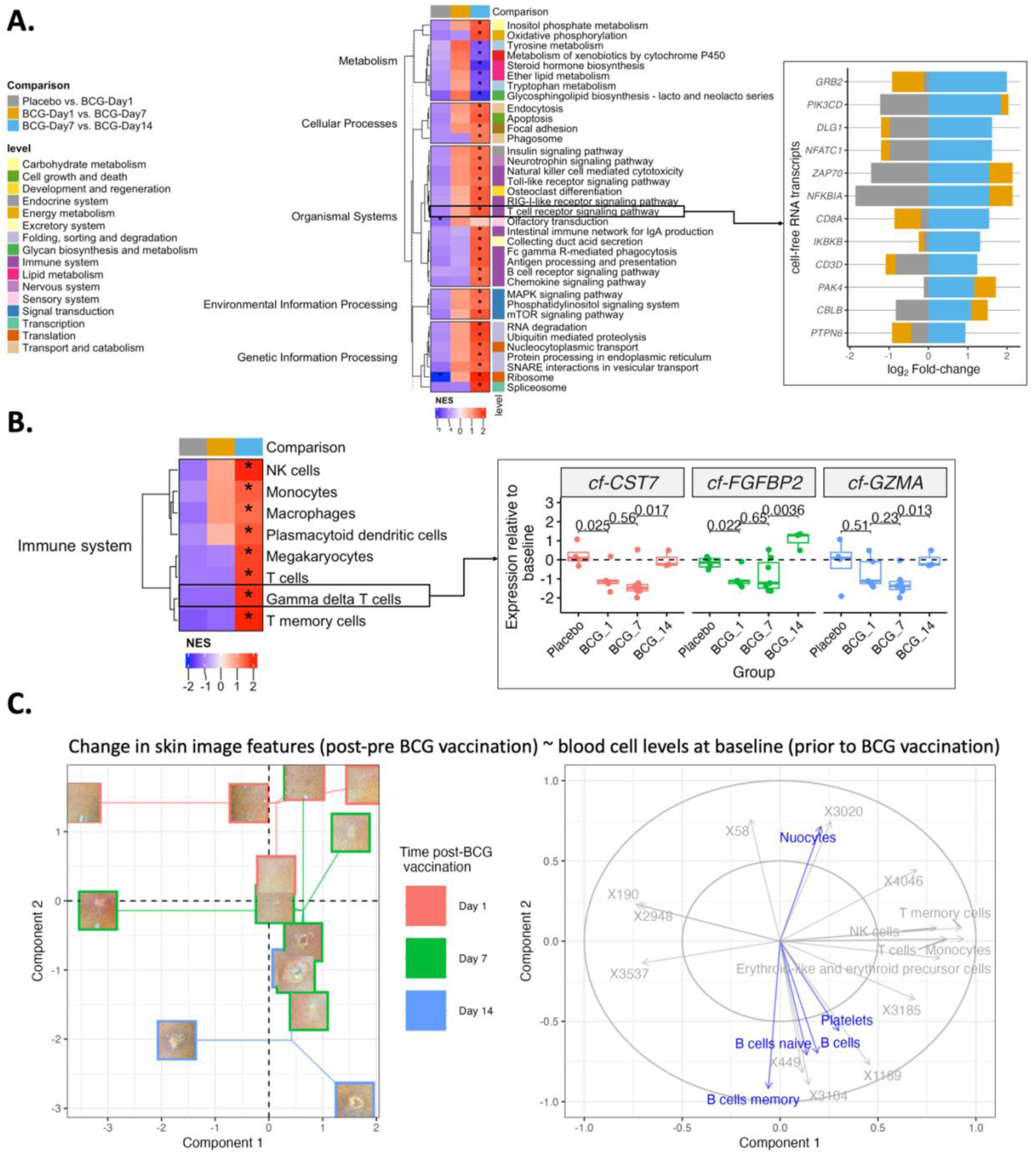
Cell-free gene-expression analysis comparing placebo and BCG samples over time in the blood. A. Heatmap of significant KEGG pathways (BH-FDR<10%) in blood based on geneset enrichment analysis of various comparisons and tissue regions. The stacked bar plot shows the fold-change of significant (*p<0.05*) gene-transcripts comparing day 7 with day 14 samples that are part of the T cell receptor signaling pathway. B. Heatmap of significant cell-types (BH-FDR<10%) in blood based on geneset enrichment analysis of various comparisons and tissue regions. Barplots for significant (*p<0.05*) gene-transcripts comparing day 7 and day 14 samples that are part of the γδ T cells geneset. C. Sparse partial least-squares (sPLS) analysis between blood cell levels (summarized from gene-expression data) at baseline (prior to BCG vaccination) and image features derived from dermatoscopic images of the site of injection (change in image features from baseline to a given day). Left: Component plot depicting the samples grouped based on the day post-BCG vaccination, aligned along the y-axis (component 2). Images are overlayed onto each sample and connected for each day using the centroid between points. Right: Correlation circle depicting the relationship between sPLS components and original image features (beginning with an X) and blood cells. Cells in blue are associated with component 2.

### BCG-induces coordinated molecular mechanisms across the skin and blood

To identify mechanisms that span the skin and blood after BCG vaccination, we integrated digital dermatoscopic images of the skin-surface with gene-expression and cell-type levels from skin and blood, using Regularized Generalized Canonical Correlation Analysis^10^ (RGCCA). RGCCA was used to maximize the correlation between differentially expressed (FDR < 10%) image features, gene-transcripts and cell levels (summarized from gene-expression data, see Methods) from skin samples, with blood cell levels (Fig.5A). RGCCA was used to extract two components that explained >55% of the variation across all datasets (Fig.S3). Component 1 discriminated placebo and BCG vaccination timepoints, whereas Component 2 separated day 14 BCG samples from the rest (Fig.5B). Fig.5C depicts the correlation between image features, gene-transcripts and cells with components 1 and 2 (top 5 features from each dataset). The dermatoscopic image features (see variables starting with an X) aligned in the direction of day 14 post-BCG samples, suggesting that these image features are representative of pustule formation in the skin. Component 1 which discriminated between placebo and BCG samples over time was positively correlated with γδ T cells, and *C1QB* in the epidermis and dendritic cells and *CERS2* in the dermis+hypodermis. The levels of these genes and cells were the lowest in placebo samples and increased in BCG samples over time (Fig.5C right). Conversely, *CLDN18*, and *RTP3* in the epidermis and *SRPK3*, and *MRGPRX1* in the dermis+hypodermis, were negatively correlated with Component 1. That is, their levels were highest in placebo samples and decreased in BCG samples over time (Fig.5C left). Component 2 (BCG response by 2 weeks) was positively correlated with γδ T cells, and platelets in blood and *SELENOP* and *PAPLN* in the dermis+hypodermis. Levels of these genes and cells were elevated at day 14 as compared to placebo and week 1 BCG samples (Fig.5C top). Conversely, *HS3ST3B1*, *PDE4D* and *TNFRSF10C* in the epidermis and *OR4F4* in the dermis+hypodermis, were negatively correlated with Component 2. That is, their levels were up-regulated at day 1 post-BCG vaccination but decreased over time, with levels returning to baseline and placebo samples at day 14 (Fig.5C bottom).

**Figure 5.**
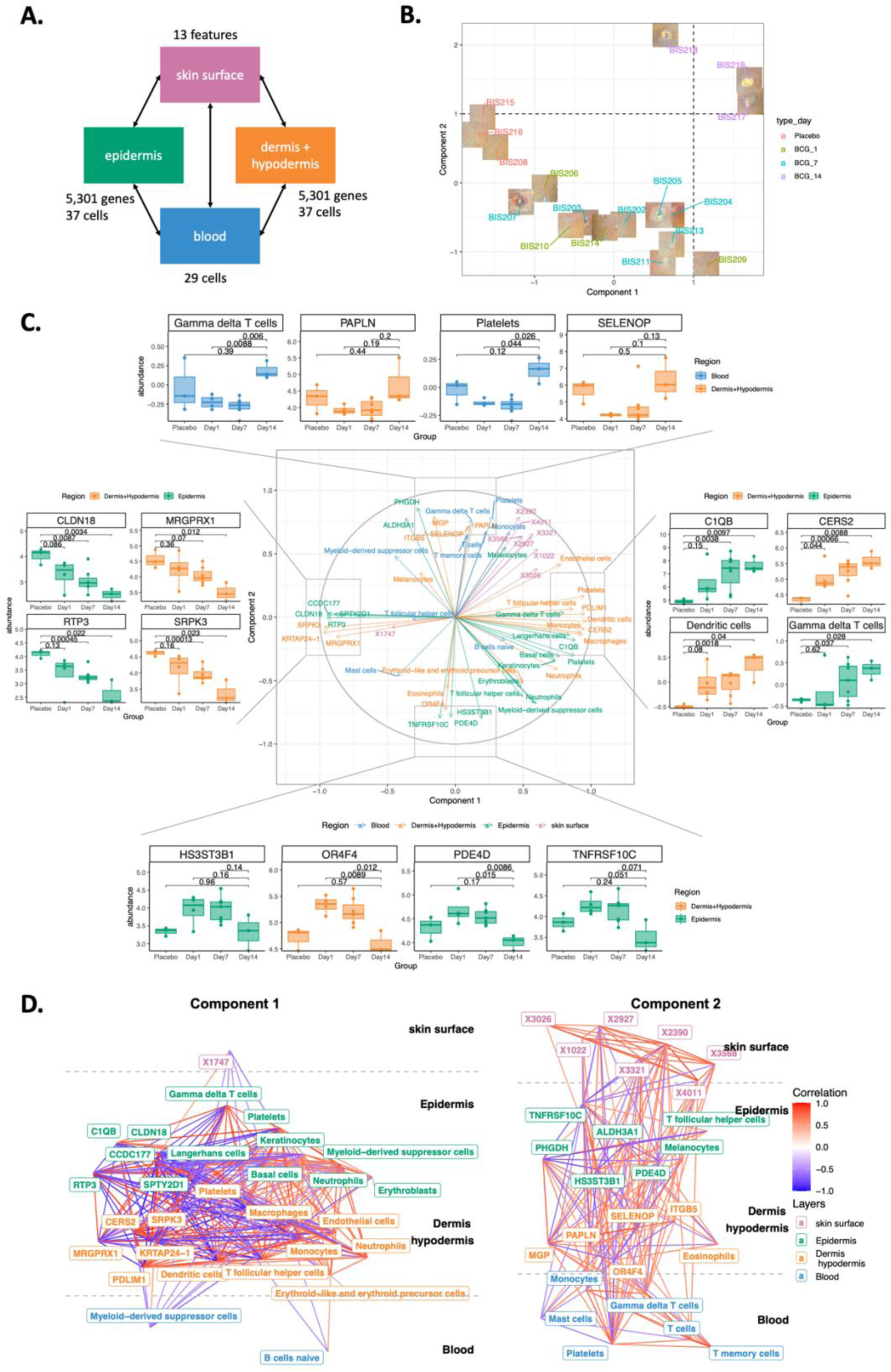
Integrating digital pathology, spatial transcriptomics of the skin and blood transcriptomics. A. Number of variables and design matrix used in the integrative analysis to maximize the covariance between skin and blood datasets. B. Dimension reduction plot using the consensus components (superblocks) across all datasets. C. Correlation circle depicting the association between the top 5 variables genes/cells identified in the integration analysis with components 1 and 2. D. Networks depicting the connections between highly correlated variables specific to component 1 and component 2 from the integrative analysis. Edges with a Pearson correlation (absolute value) greater than 0.5 are shown.

Fig.5D depicts hierarchical networks across the skin and blood for component 1 in which features are different between placebo and BCG samples over time (left) and component 2 in which features are different between all samples and day 14 samples (right). Component 1 may represent a response that is sustained in the skin throughout the two-week study period. Most connections in this network are between gene-transcripts and cells in different regions of the skin. Component 2 may indicate a secondary response that span both skin and blood, comprised several dermatoscopic features and blood cells. Despite the inherent complexity, these clear connections indicated a coordinated molecular response across the skin and blood after BCG vaccination related to the macroscopic changes observed in the skin.

## Discussion

This data represents the first holistic view of the human body’s acute response to BCG during the first two weeks post-vaccination in a medium to high TB risk area (West Africa). By investigating both local and systemic responses to vaccination we were able to identify novel molecular responses. This data also reveals biomarkers that in the future may allow prediction of vaccine efficacy, crucial for the understanding of BCG’s immune effects and the acceleration of TB vaccine development. Each of our data modalities consisted of features that were significantly altered after BCG vaccination, including dermatoscopic changes of the skin (indicating skin reaction formation) as well as changes in structural features of the skin (indicating changes in cell morphology), reflecting molecular changes in gene-and cell-expression across layers of the skin and peripheral blood over time. Integration of these dynamic changes across space and time enabled identification of variables (components) that likely represent molecular mechanisms (spanning multiple modalities) induced by BCG vaccination.

Mycobacteria introduced through BCG vaccine administration could be identified in the skin for the first week post-vaccination, but not later when the development of a pustule was well underway. This indicates that the development of the later skin responses and subsequent scar formation were not driven by local presence of mycobacteria, but rather the immunological cascade initiated by their initial presence. Dermatoscopic changes observed in the skin images such as hyperpigmentation were corroborated by the gene-expression data through increased melanocytes at later timepoints. Further, a reduction in erythema was observed over time and this aligned with the significant increase in erythroblasts as well as erythroid-like and erythroid precursor cells at day 1 in the epidermis, and a decrease of the same at the later time points.

We found that BCG induced dramatic changes in cellular composition of the skin within a day after vaccination involving resident macrophages and dendritic cells. We also identified rapid NK cell activation which may indicate the presence of tissue-resident or rapidly invading blood-derived NK cells^11^. These local responses in the skin differed by skin layer. Initially most cellular responses were observed in the epidermis, the site where BCG was injected. However, after the first day, significant changes also occurred in the dermis+hypodermis, which then persisted over two weeks, suggesting the initial epidermal response had moved to deeper skin layers. Cell types that were found to be up-regulated subsequently included fibroblasts and platelets which are involved in wound healing, indicating that mechanisms associated with eventual scar formation appear to be activated early^12,13^. Unlike the local skin response to BCG, which was evident already on day 1, the systemic response (as assessed by cell-free plasma transcripts) was only observed by two weeks after vaccination. Systemic increased activity in cell types identified via cell-free RNA during the second week not only included NK cells, but also monocytes, macrophages, plasmacytoid dendritic cells, megakaryocytes, T cells, γδ T cells and T memory cells. These cell-types are known to respond to TB infection, through ingestion of bacteria^14^ (monocytes, macrophages), killing infected cells^15^ (NK cells and γδ T cells) and developing long-term immunity^16^ (T memory cells). Our results also indicated that higher levels of B cells and platelets and lower levels of nuocytes were associated with particular macroscopic changes in the skin. For example, introducing large number of B cells into the skin can promote tissue regeneration and accelerate would healing^17^. Nuocytes are Type 2 innate lymphoid cells known to respond to helminth infection^9^, such that reduced number of nuocytes may indicate a reduced capacity to clear helminth infection. Importantly, since levels of these cell types were inferred based on cfRNA transcripts that are hypothesized to be derived from tissues, our data may indicate expansion of these cell types within host tissues and increased circulating levels of these cells *per se*.

Analyzing each modality separately was useful in identifying genes and cells associated with BCG vaccination. However, the multimodal integration across space and time revealed two prominent patterns of BCG across the skin and blood. The first pattern consisted of gene-transcripts and cell-types in the skin such as *CERS2, PDLIM1, C1QB,* neutrophils, platelets, and γδ T cells whose levels were different between placebo and post-BCG samples over time and inversely associated with myeloid-derived suppressor cells (MDSCs) in the blood. Further, MDSCs in the epidermis were found to be negatively correlated with MDSCs in the blood, possibly due to trafficking of these cells into the skin such that a reduction of MDSCs in the blood is caused by increased MDSC levels in the skin. The presence of MDSCs in the skin suggests a counterbalancing role to γδ T cells which promote inflammation to fight bacterial infection. Complement Component 1, Q Subcomponent, B Chain (*C1QB*) in the epidermis was elevated in BCG samples compared to placebo and increased over time. Further, *C1QB* levels were positively correlated with monocytes and macrophages in the dermis+hypodermis. C1q has been shown to be produced by monocytes and may serve as a biomarker to distinguish active from latent TB^18^.

The second prominent pattern centred around T memory cells, γδ T cells, and platelets in the blood and *PAPLN,* and *SELENOP* in the skin, whose levels increased in the second week post-BCG vaccination as compared to the first week. Memory and γδ T cells in peripheral blood are recruited to sites of inflammation in response to infection^19,20^, however since this response occurred during the second week post-BCG vaccination, it is likely secondary to the initial immune response in the skin at day 1. The response observed in the skin at two weeks post-BCG vaccination included up-regulation of *PAPLN* genes involved in extracellular matrix organization, which may facilitate immune cell infiltration. At the same time, the antioxidant genes such as *SELENOP* were increased at two weeks, which may be a protective response to prevent host tissue damage^21^. Both of these genes were positively associated with γδ T cells and platelets in the blood.

An integration analysis also revealed gene and cellular patterns in the blood that were associated with macroscopic changes observed by dermatoscopic images at day 14. As such, the dermatoscopic local skin response to BCG appears to be mirrored by a systemic immune response. A previous study showed that baseline production of IFN-γ was associated with BCG vaccination scar size^22^. This may suggest that blood biomarkers may be used to predict progression of the local skin response and possibly subsequent scar formation after BCG vaccination.

We are of course limited by a relatively small sample size, albeit with unprecedented deep molecular phenotyping of each participant. Second, the study’s time frame was confined to early molecular changes observed up to 14 days post-BCG vaccination. While this allowed us to capture immediate and short-term responses, it does not provide insights into long-term immune mechanisms or initiation of any protective effects. Our results are likewise confined to a specific global region and adult women only, meaning that environmental and genetic factors unique to this population as well as age or sex could influence the immune responses observed. However, our data show that it is possible to conduct deep molecular phenotyping in a low resource setting and that this approach can reveal the complex molecular pathways initiated by BCG.

In summary, this data represents the first insights into the dynamic molecular mechanisms underlying the acute host response to BCG vaccination spanning time and space. This includes both local intercellular communication exhibiting complex waves over time (days) and space (layers of the skin), as well as perturbations of systemic signals detected by the cell-free blood-based transcripts. Our findings both confirm known biological evidence and generate new hypotheses. While these novel findings require validation, our data highlight that multimodal spatiotemporal integration can reveal key interactions across space and time. This holistic assessment of the host response to BCG along with the possibility of non-invasive sampling (dermatoscopy) predicting outcome at time of vaccination could help accelerate of vaccine development in the fight against TB.

## Methods

### Study design

The study was approved by the National Committee for Ethics in Health Research of Guinea-Bissau (CNES), reference number 027/CNES/INASA/2023. We performed this experimental medicine study of 19 adult women (HIV-negative, nonpregnant, BCG-scar-negative and without signs of active TB) who participated in this study after written informed consent. Of the 19 participants, 15 received BCG and the remaining 4 (randomized to placebo) were offered BCG at the end of the trial. Participants were recruited in Bandim Health Project’s urban study area. The Bandim Health Project’s (BHP, www.bandim.org) Health and Demographic Surveillance System (HDSS) site follows a population of 102,000 individuals in six suburbs in Bissau, Guinea-Bissau, with home visits and through surveillance at the nearby national hospital, and has a long track record of conducting large-scale RCTs of different vaccines with overall and infectious disease morbidity and mortality as outcomes. Females above 18 were invited to participate in the study. All participants were screened for HIV via a rapid antigen blood test and for TB via a previously validated TB score [https://pubmed.ncbi.nlm.nih.gov/23113626/] based on physical exam and medical history performed by the study physician. Pregnancy was assessed via a rapid human choriogonadotropin urine test. Presence of a BCG scar was assessed through inspection of the deltoid area of both arms. Participants who were negative for HIV and TB and who were not pregnant and who did not have a BCG scar were invited to participate in the study. In a subsequent study using the same methodology, we recruited participants that had a BCG scar (data currently being analyzed).

### Intervention and randomization

Participants were randomized to BCG vs. Saline and to a follow-up day of 1, 7, or 14 days after the baseline visit. Participants received a standard 1.0 mL dose of BCG vaccine (AJ Vaccines, Denmark, Denmark Strain of BCG) or 0.9% injection-grade saline for placebo controls. Immediately following the intervention, the height and width of the injection bleb were measured with a ruler to document the inoculum size. We employed a dermatoscope (Dermlite DL) to collect high-quality standardized images of the skin during the baseline visit (to photograph the injection bleb) and during the follow up visit, to photograph the skin reaction to the intervention prior to collecting skin biopsy samples.

### Sample collection

Blood samples were collected in 4.0 mL K2EDTA tubes from each participant at baseline (prior to vaccination) and at follow-up (one of days 1, 7 or 14). Since HIV tests were performed immediately after blood draw, processing did not commence until participants were confirmed HIV negative, 15-20 minutes following blood sample collection. Whole blood was preserved at the site of blood draw by placing 1.3 mL whole blood into *247* µL cfRNA preservative (Norgen Cat. 63950) using DNase/RNase DNA Lo-Bind tubes and mixed gently using DNase/RNase-free wide-bore pipette tips. Blood samples were then transported to the laboratory for further processing, where the cfRNA preservative-blood mixture was centrifuged at 460 x g for 20 minutes and the top 90% of the plasma lifted to a new 2.0 mL tube. This tube was spun again at 2600 x g for 15 minutes to yield cell free plasma. Taking care to avoid the pellet, the plasma was transferred to a new 2.0 mL microtube and stored at -80 ℃.

Each participant donated one skin punch biopsy over the course of the study, (i.e., day 1, 7, 14 post-BCG). Biopsy samples were collected after blood draw. The previous BCG or placebo administration site was identified by scar formation and/or from photographs taken at the time of administration. The biopsy site was marked with a skin marker for those participants with no scar visible. Skin was anaesthetised by application of an ice pack for 5-10 minutes and then cleaned with a regular alcohol wipe as is standard before a blood draw. The sterile, single-use skin punch was opened and a single, 2 mm biopsy was obtained. The skin samples were processed on site, *i.e.* immediately, stored in 1.0 mL paraformaldehyde and stored at room temperature for 24 hours. The sample was then rinsed with 10 mL sterile PBS and frozen at -80°C in an RNAase inhibitor (RNALater, ThermoFisher).

### Tissue sectioning and staining

Sequential five-micron tissue sections were cut in quadruplicate from archived FFPE-embedded specimens onto charged slides. Two slides were stained: one with hematoxylin and eosin (H&E) and the other with Ziehl–Neelsen^23^ (ZN), following established protocols. Stained slides were digitally scanned for evaluation by a pathologist. Of the remaining three slides, one was used for optimization profiling, while the other two were utilized for transcriptomic profiling using the GeoMx™ DSP.

### GeoMx™ transcriptome DSP

Spatial profiling for the transcriptomic analyses was undertaken as recommended by NanoString Technologies (Seattle, Washington, USA). Sectioned tissue was baked in a drying oven for one hour at 60⁰C and then underwent antigen retrieval followed by treatment with Proteinase K to expose RNA targets. Slides for profiling of the skin biopsy were incubated with the Human Whole Transcriptome Atlas according to the manufacturer’s instructions (MAN-10150-04). Two non-segmented ROIs and two segmented ROIs were collected from epidermis and dermis+hypodermis areas of each sample guided by a pathologist, avoiding hemorrhagic and acellular portions of tissues. ROIs were matched as best as possible between serial sections for RNA profiling and between study visits. Following collection of the indexing oligos into a 96 well plate, samples were sequenced on a NovaSeq system (Illumina) at the Australian Genome Research Facility (AGRF) according to Nanostring recommendations.

### Cell-free RNA

The 38 human plasma samples were shipped to Norgen Biotek Corporation for cell-free RNA sequencing. Norgen’s Plasma/Serum RNA purification Midi Kit (# 56100) was used to extract RNA. The Ilumina NextSeq 500 was used to perform RNA sequencing. Adapter trimming was performed using Cut-Adapt, alignment was performed using the START aligner (UCSC hg38 Human reference) and the count matrix was generated using FeatureCounts. The count data was normalized to log_2_ counts per million: 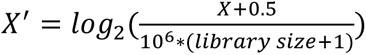, where library size is the total counts of all genes for a given sample.

### Image processing

Dermatoscopic images of the site of injection and immunofluorescence images of the regions of interested were cropped to remove background and rescaled to images of 224x224 pixels. After normalization, feature extraction was performed by applying the AlexNet per-trained computer vision model, resulting in a 4,096 feature-vector for each image (using the Python libraries torchvision v0.19 and PyTorch v2.2.2). Image features with zero variance were removed prior to downstream analysis.

### Differential expression analysis

Three comparisons were performed for each dataset: Placebo *vs.* BCG at day 1, BCG at day 1 *vs.* BCG at day 7 and BCG at day 7 *vs.* BCG at day 14. All datasets consisted of 19 samples, except the dematoscopic image data and cell-free RNA data which consisted of 38 samples (pre and post vaccination). For each patient, pre-expression was subtracted from post-expression resulting in a dataset for the 19 participants [Placebo (*N*=4), BCG day 1 (*N*=5), BCG day 7 (*N*=7) and BCG day 14 (*N*=3)]. Moderated t-test using the Linear Models for Microarrays and RNA-seq (limma R-library, v3.58.1). The Benjamini-Hochberg False Discovery Rate (BH-FDR) was used to correct for multiple testing and threshold of 10% was used to assess statistical significance.

### Gene set enrichment analysis

The test-statistics estimated using the differential expression analysis were used to perform gene set enrichment analysis (GSEA) using fgsea R-library (v1.30.0). Gene sets included The Kyoto Encyclopedia of Genes and Genomes (KEGG). Cell-type markers genes were download from PanglaoDB.

### Gene set variation analysis

Gene set variation analysis (GSVA) was used to summarize the epidermis and dermis+hypodermis specific spatial-gene expression data using marker genes for significant marker genes only. GSVA was used to summarize the cell-free RNA data using marker genes for blood and imme cells from PanglaoDB. The resulting datasets were used as inputs to the integration analyses.

### Integration analyses

Sparse Partial Squares (mixOmics R-library v6.28.0) was used to regress the change in image features (after BCG vaccination minus baseline) onto cell-type levels at baseline (prior to BCG vaccination). Two components were retained and 5 variables were selected for each component.

Regularized Generalized Canonical Correlation Analysis (RGCCA R-library v3.0.3) was used to integrate across the skin surface, skin layers and blood (6 datasets). The skin surface datasets consisted of image features extracted from dermatoscopic images. The 4 datasets related to the skin layers consisted of both gene-expression and cell-types (summarized using gene-expression) from epidermis and dermis+hypodermis regions. The blood datasets consisted of only cell-types (summarized from plasma cell-free RNA). For all datasets, only differentially expressed variables in at least one of the comparisons (Placebo *vs.* day 1, day 1 *vs.* day7 and day 7 *vs.* day 14) were used in the integration analysis. RGCCA was used to integrate these six datasets by maximising the sum of the pairwise correlations between the skin and blood datasets only, and two components were retained.

## Data availability

All data will be made publicly available upon acceptance of the manuscript. Specifically, we will submit the transcriptomics data presented in this publication to the NCBI Gene Expression Omnibus and provide the respective accession numbers. Other data sets will be archived on ImmPort, again with the respective accession numbers provided.

## Data availability

All data files associated with these analyses will be uploaded to Zenodo at the time of publication.

## Code availability

All custom code developed for these analyses have been deposited on GitHub: https://github.com/CompBio-Lab/bcg_skinblood_crosstalk. R version and associated library names and version can be found in https://github.com/CompBio-Lab/bcg_skinblood_crosstalk/blob/main/renv.lock. Python version and associated library names and versions can be found in https://github.com/CompBio-Lab/bcg_skinblood_crosstalk/blob/main/scripts/bcg_dermatoscope.yml.

## Notes

### Competing Interest Statement

The authors have declared no competing interest.

## References

1. Global Tuberculosis Report 2024. https://www.who.int/teams/global-tuberculosis-programme/tb-reports/global-tuberculosis-report-2024.

2. Bæk, O. et al. The mark of success: The role of vaccine-induced skin scar formation for BCG and smallpox vaccine-associated clinical benefits. Semin Immunopathol 46, 13 (2024).

3. Minassian, A. M. et al. A Human Challenge Model for Mycobacterium tuberculosis Using Mycobacterium bovis Bacille Calmette-Guérin. The Journal of Infectious Diseases 205, 1035–1042 (2012).

4. Harris, S. A. et al. Evaluation of a Human BCG Challenge Model to Assess Antimycobacterial Immunity Induced by BCG and a Candidate Tuberculosis Vaccine, MVA85A, Alone and in Combination. Journal of Infectious Diseases 209, 1259–1268 (2014).

5. Minhinnick, A. et al. Optimization of a Human Bacille Calmette-Guérin Challenge Model: A Tool to Evaluate Antimycobacterial Immunity. J Infect Dis. 213, 824–830 (2016).

6. Alorainy, M. et al. A Systematic Review of Diagnoses With Rosettes Under Dermoscopy. Dermatology Practical & Conceptual 14, e2024125 (2024).

7. Haspeslagh, M. et al. Rosettes and other white shiny structures in polarized dermoscopy: histological correlate and optical explanation. Acad Dermatol Venereol 30, 311–313 (2016).

8. Franzén, O., Gan, L.-M. & Björkegren, J. L. M. PanglaoDB: a web server for exploration of mouse and human single-cell RNA sequencing data. Database 2019, (2019).

9. Neill, D. R. et al. Nuocytes represent a new innate effector leukocyte that mediates type-2 immunity. Nature 464, 1367–1370 (2010).

10. Tenenhaus, A. & Tenenhaus, M. Regularized generalized canonical correlation analysis for multiblock or multigroup data analysis. European Journal of Operational Research 238, 391–403 (2014).

11. Narni-Mancinelli, E., Berruyer, C. & Vivier, E. On blood and tissue-resident natural killer cells. Immunity 57, 6–8 (2024).

12. Cialdai, F., Risaliti, C. & Monici, M. Role of fibroblasts in wound healing and tissue remodeling on Earth and in space. Front Bioeng Biotechnol 10, 958381 (2022).

13. Thomas, H. M., Cowin, A. J. & Mills, S. J. The Importance of Pericytes in Healing: Wounds and other Pathologies. Int J Mol Sci 18, 1129 (2017).

14. Pahari, S. et al. Reinforcing the Functionality of Mononuclear Phagocyte System to Control Tuberculosis. Front Immunol 9, 193 (2018).

15. Chowdhury, R. R., et al. NK-like CD8+ γδ T cells are expanded in persistent Mycobacterium tuberculosis infection and chronic inflammation. Sci Immunol 8, eade3525 (2023).

16. Counoupas, C. & Triccas, J. A. The generation of T-cell memory to protect against tuberculosis. Immunol Cell Biol 97, 656–663 (2019).

17. Sîrbulescu, R. F. et al. Mature B cells accelerate wound healing after acute and chronic diabetic skin lesions. Wound Repair Regen 25, 774–791 (2017).

18. Cai, Y. et al. Increased complement C1q level marks active disease in human tuberculosis. PLoS One 9, e92340 (2014).

19. Gray, J. I., Westerhof, L. M. & MacLeod, M. K. L. The roles of resident, central and effector memory CD4 T-cells in protective immunity following infection or vaccination. Immunology 154, 574–581 (2018).

20. Nanda, N. & Alphonse, M. P. From Host Defense to Metabolic Signatures: Unveiling the Role of γδ T Cells in Bacterial Infections. Biomolecules 14, 225 (2024).

21. Zhang, J., Saad, R., Taylor, E. W. & Rayman, M. P. Selenium and selenoproteins in viral infection with potential relevance to COVID-19. Redox Biol 37, 101715 (2020).

22. Moorlag, S. J. C. F. M. et al. Multi-omics analysis of innate and adaptive responses to BCG vaccination reveals epigenetic cell states that predict trained immunity. Immunity 57, 171–187.e14 (2024).

23. Vilchèze, C. & Kremer, L. Acid-Fast Positive and Acid-Fast Negative Mycobacterium tuberculosis: The Koch Paradox. Microbiol Spectr 5, 10.1128/microbiolspec.tbtb2-0003#2015.

