## Supplement for "Spatiotemporal-multimodal integration reveals BCG-induced skin-blood crosstalk"

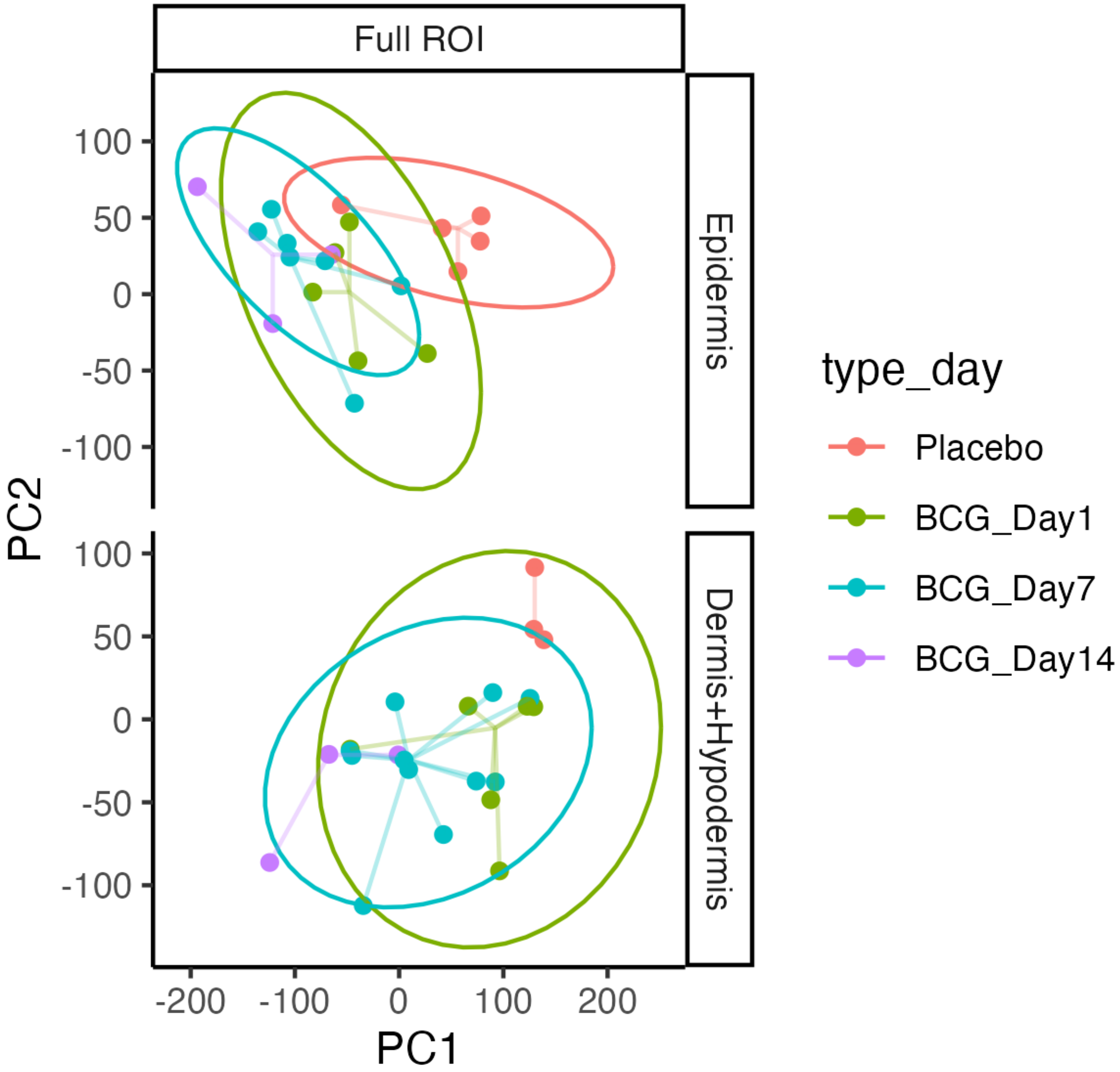


Figure S1. Principal component analysis of spatial gene-expression data for the epidermis and combined dermis+hypodermis.

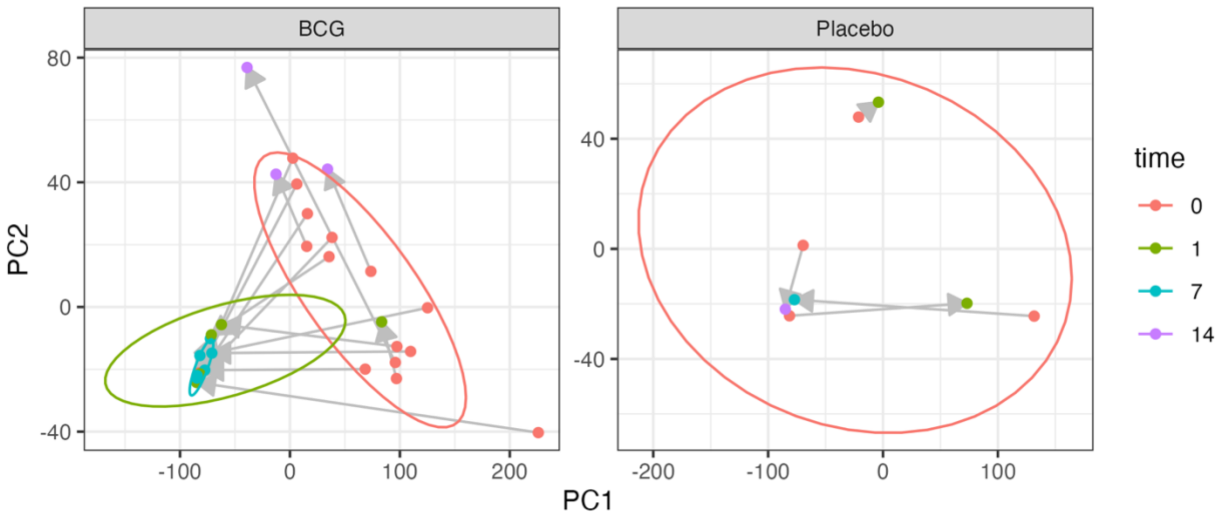


Figure S2. Principal Component Analysis of blood gene-expression data for all 38 cell-free RNA samples

Figure S3. Variation explained using RGCCA.

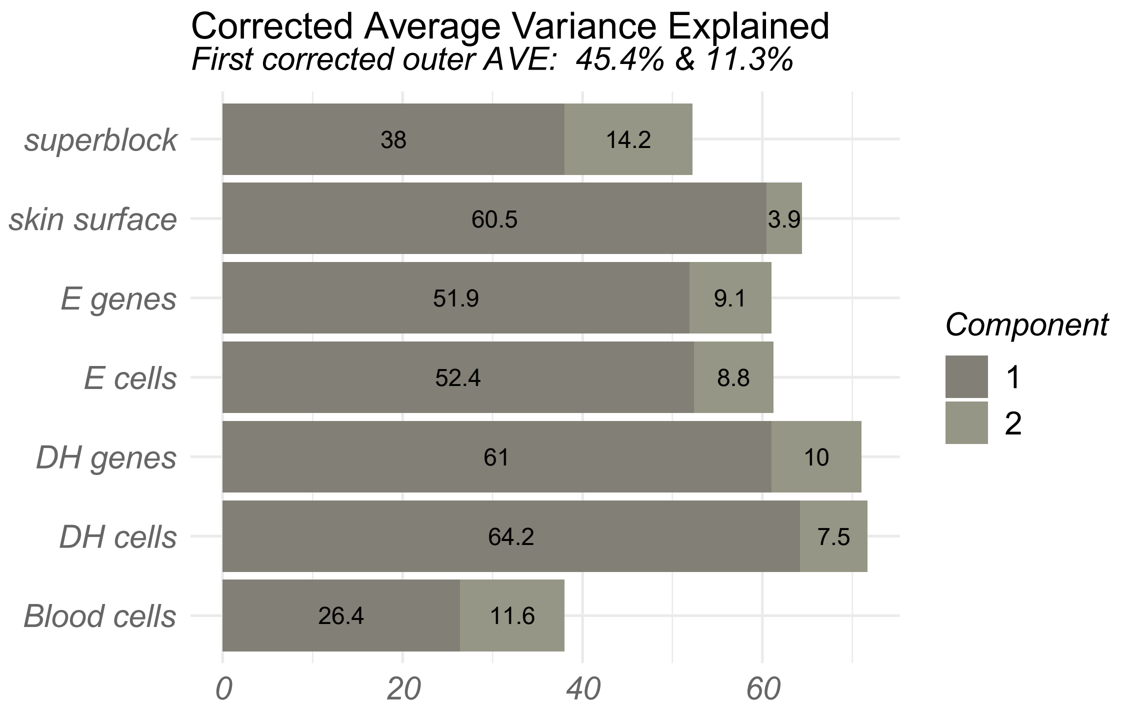


**Table S1: Dermatoscopic images of skin with either placebo or BCG vaccination (before and after).**

Dermatoscopic images

**Placebo**

| Baseline | Day 1 |
|---|---|

| | |
|---|---|
| SITU018_BIS018 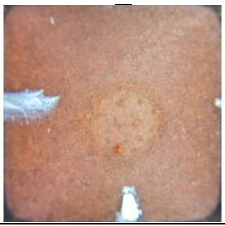 | SITU018_BIS215 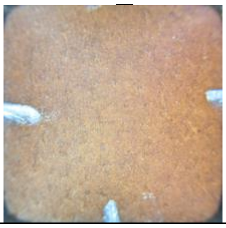 |
| SITU019_BIS019 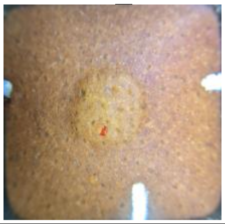 | SITU019_BIS216 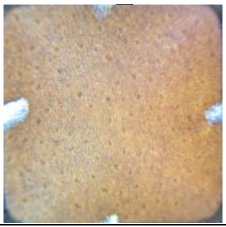 |

| Baseline | Day 7 |
|---|---|
| SITU009_BIS009 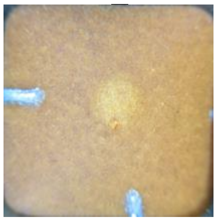 | SITU009_BIS208 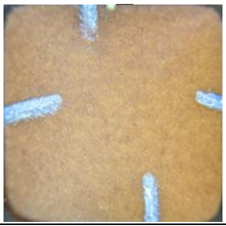 |

| Baseline | Day 14 |
|---|---|
| SITU001_BIS001 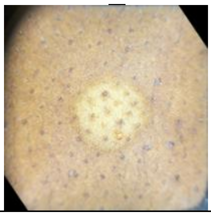 | SITU001_BIS212 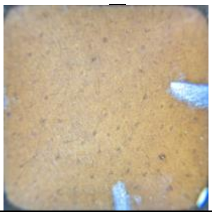 |

**BCG**

| Baseline | Day 1 |
|---|---|
| SITU011_BIS011 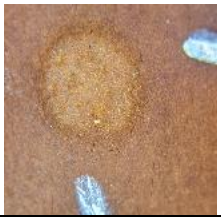 | SITU011_BIS201 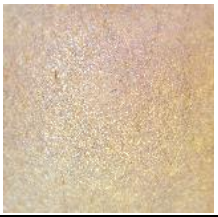 |
| SITU014_BIS014 | SITU014_BIS206 |

| | |
|---|---|
| 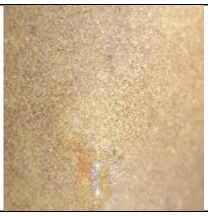 | 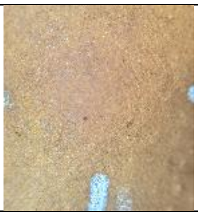 |
| SITU015_BIS015 | SITU015_BIS209 |
| 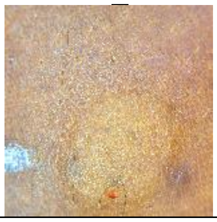 | 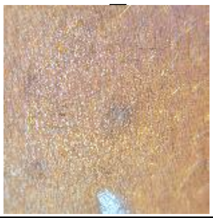 |
| SITU016_BIS016 | SITU016_BIS214 |
| 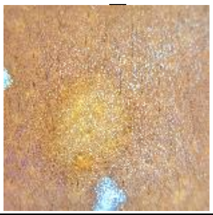 | 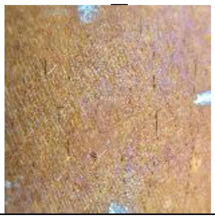 |
| SITU017_BIS017 | SITU017_BIS210 |
| 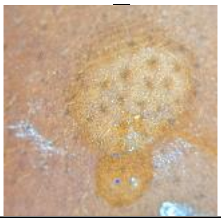 | 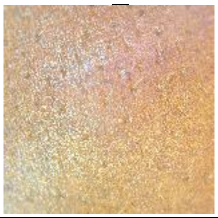 |

| Baseline | Day 7 |
|---|---|
| SITU005_BIS005 | SITU005_BIS202 |
| 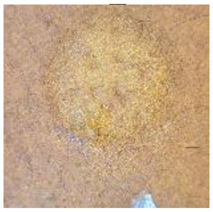 | 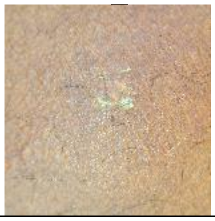 |
| SITU006_BIS006 | SITU006_BIS203 |
| 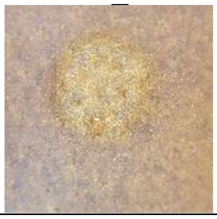 | 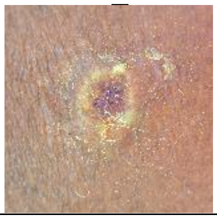 |
| SITU007_BIS007 | SITU007_BIS204 |

| | |
|---|---|
| 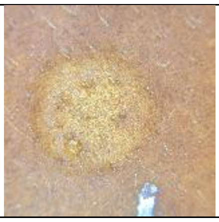 | 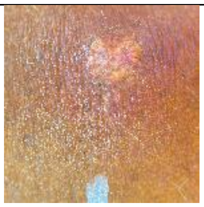 |
| SITU008_BIS008 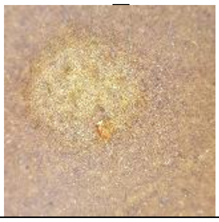 | SITU008_BIS205 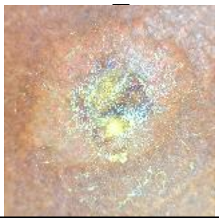 |
| SITU010_BIS010 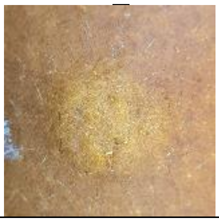 | SITU010_BIS207 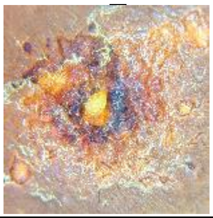 |
| SITU012_BIS012 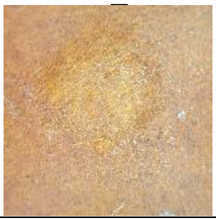 | SITU012_BIS211 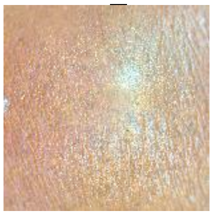 |
| SITU013_BIS013 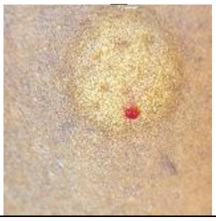 | SITU013_BIS213 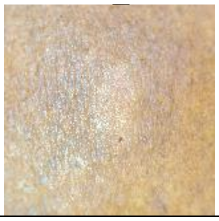 |

| Baseline | Day 14 |
|---|---|
| SITU002_BIS004 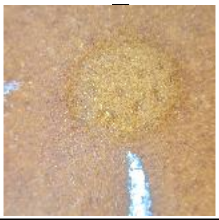 | SITU002_BIS217 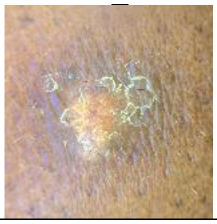 |
| SITU003_BIS002 | SITU003_BIS218 |

| | |
|---|---|
| 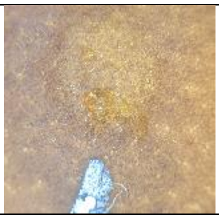 | 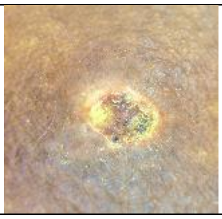 |
| SITU004_BIS003 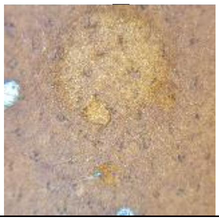 | SITU004_BIS219 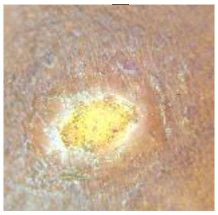 |
